# The Role of Recombination in Evolutionary Rescue

**DOI:** 10.1101/022020

**Authors:** Hildegard Uecker, Joachim Hermisson

## Abstract

How likely is it that a population escapes extinction through adaptive evolution? The answer to this question is of great relevance in conservation biology, where we aim at species’ rescue and the maintenance of biodiversity, and in agriculture and epidemiology, where we seek to hamper the emergence of pesticide or drug resistance. By reshuffling the genome, recombination has two antagonistic effects on the probability of evolutionary rescue: it generates and it breaks up favorable gene combinations. Which of the two effects prevails, depends on the fitness effects of mutations and on the impact of stochasticity on the allele frequencies. In this paper, we analyze a mathematical model for rescue after a sudden environmental change when adaptation is contingent on mutations at two loci. The analysis reveals a complex nonlinear dependence of population survival on recombination. We moreover find that, counterintuitively, a fast eradication of the wildtype can promote rescue in the presence of recombination. The model also shows that two-step rescue is not unlikely to happen and can even be more likely than single-step rescue (where adaptation relies on a single mutation), depending on the circumstances.

## Introduction

Populations facing severe environmental change need to adapt rapidly to the new conditions, or they will go extinct. The most prominent examples for evolutionary rescue in natural populations are provided by failed eradication of pathogens or pests that develop resistance against drugs or pesticides. Understanding which factors drive the evolution of resistance has been a concern since the application of drugs and pesticides. In recent years, the topic of evolutionary rescue has attracted increasing interest of evolutionary biologists at a broader front. Both theoretical models and laboratory experiments have been used to investigate the influence of many genetic or environmental factors on the survival probability of an endangered population, e.g., the importance of standing genetic variation, sexual reproduction, the history of stress, the severity and speed of environmental deterioration, or population structure (Bell and Collins (2008); Bell and Gonzalez (2009, 2011); Orr and Unckless (2008, 2014); Agashe *et al.* (2011); Lachapelle and Bell (2012); Gonzalez and Bell (2013); Uecker *et al.* (2014); see also the reviews by Alexander *et al.* (2014) and Carlson *et al.* (2014)). Despite significant progress, a largely open area in research on rescue concerns the influence of recombination on the probability of population survival.

Recombination has two fundamental effects on adaptation that work against each other: it brings favorable gene combinations together but it also breaks them up. Recombination hence has the potential both to promote rescue or to impede it. In classical population genetics (assuming a constant population size), the interplay of the two opposing effects of recombination has been an intensively studied problem for decades (e.g., reviewed in Barton and Charlesworth, 1998; Otto, 2009; Hartfield and Keightley, 2012). Fundamentally, recombination acts to reduce the linkage disequilibrium between alleles. For two loci with two alleles each, recombination increases the number of double mutants if the linkage disequilibrium (LD) between the mutant alleles is negative; it decreases them when LD is positive; and it has no effect if the loci are in linkage equilibrium. A major source of linkage disequilibria is epistasis with negative epistasis leading to negative LD and positive epistasis leading to positive LD (Felsenstein, 1965; Kouyos *et al.*, 2007). For evolutionary rescue, the shift of the environment from original to perturbed conditions may lead to a change in epistasis during the course of evolution, which adds new aspects to the problem of how recombination affects adaptation.

Combination drug therapy (as well as the use of herbicide mixtures in agriculture) seeks to limit the evolution of resistance by increasing the number of mutations that are required to restore fitness above one. Understanding under which conditions recombination can undermine this strategy is of great relevance to plan a successful treatment. Consequently, the effect of recombination on the evolution of drug resistance has attracted considerable attention from epidemiologists, in particular with respect to resistance in HIV (Bretscher *et al.*, 2004; Fraser, 2005; Carvajal-RodrÍguez *et al.*, 2007; Kouyos *et al.*, 2009). However, most epidemiological models are deterministic and focus on the time to resistance rather than on the probability of resistance (Bretscher *et al.*, 2004; Fraser, 2005). An exception is the simulation study by Kouyos *et al.* (2009), which incorporates stochasticity and allows populations to go extinct. Their study demonstrates how the population dynamics affects the emergence of linkage dis-equilibria and hence the influence of recombination on the probability and time to resistance, finding that recombination usually slows down the evolution of resistance. However, the model is specific to the complex epidemiological dynamics of HIV and cannot be used to draw general conclusions about the role of recombination in evolutionary rescue.

In a general evolutionary context, theoretical models of rescue where population genetics and population dynamics are intertwined have mainly followed two routes. In one class of models, adaptation relies on changes at just a single locus, and recombination consequently does not appear (Gomulkiewicz and Holt, 1995; Iwasa *et al.*, 2003, 2004; Bell and Collins, 2008; Orr and Unckless, 2008, 2014; Martin *et al.*, 2013; Uecker *et al.*, 2014). The second class of models, motivated by conservation biology, is based on a quantitative genetics approach where (infinitely) many loci of small effect determine the fitness of an organism (Pease *et al.*, 1989; Lynch *et al.*, 1991; Lande and Shannon, 1996; Bürger and Lynch, 1995; Polechova *et al.*, 2009; Duputié *et al.*, 2012). These models usually do not incorporate an explicit genetic architecture that would allow for the investigation of the effect of recombination or assume linkage equilibrium. In their review, Carlson *et al.* (2014) refer to a single study – Schiffers *et al.* (2013) – for the effect of linkage on the probability of rescue. In their simulation study of an explicit multi-locus model, Schiffers *et al.* (2013) compare rescue probabilities for the two extreme cases of complete linkage and free recombination. In contrast to Kouyos *et al.* (2009), they find that linkage significantly decreases the probability of rescue. However, the study omits the range of intermediate linkage, and moreover, the model is taylored to consider a highly specific ecological situation of climate change in a spatially structured environment. Just as the study by Kouyos *et al.* (2009), it is not designed to serve as a baseline model.

In this paper, we set up and analyze a generic two-locus model for the role of recombination in evolutionary rescue. A population experiences a sudden severe environmental change; adaptation relies on two mutations and can both happen either from the standing genetic variation or from de-novo mutations. There are hence two phases – the time before and the time after the environmental change – during which recombination acts to increase or decrease the chances of population survival, depending on the fitness scheme and the strength of drift. We provide an accurate analytical framework based on branching process theory, complemented by computer simulations, to obtain an intuitive understanding of the principles underlying rescue under these conditions. We conclude with two notable observations that might contradict spontaneous intuition and that could be of practical relevance.

## The model

Consider a panmictic population of variable size *N* = *N* (*t*) that faces the risk of extinction after a sudden environmental change. Individuals are haploid during their selective phase. Their fitness before and after the change depends on two loci with two alleles each such that there are four genotypes: the wildtype *ab*, single mutant types *Ab* and *aB*, and the double mutant *rescue type AB*. Each generation, each individual produces a large number *X* of gametes. Mutations happen with probability *u* at each locus and in both directions. Gametes form diploid zygotes, which produce haploid offspring. The recombination probability between the two loci is *r*. Thereafter, selection takes place. By *n_ab_*, *n_Ab_*, *n_aB_*, and *n_AB_*, we denote the number of the respective genotypes in the population; hence *N* = *n_ab_* + *n_Ab_* + *n_aB_* + *n_AB_*. We obtain for the number of haploids after reproduction but before selection:

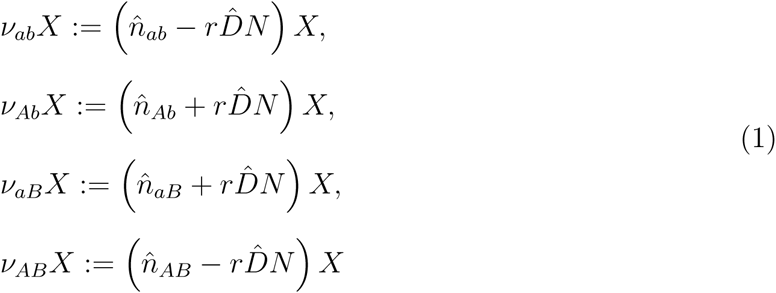

with the proportion of each genotype after mutation but before recombination

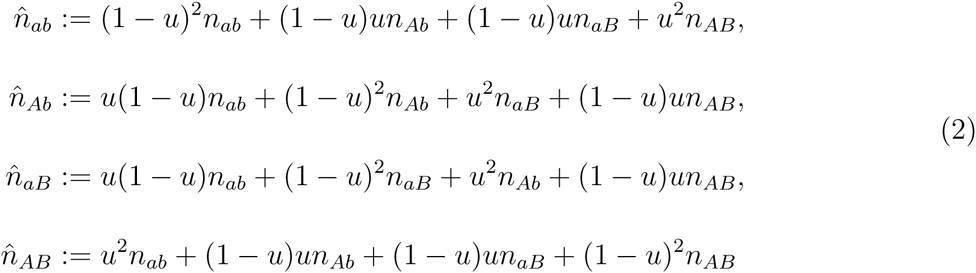

and the linkage disequilibrium (after mutation)

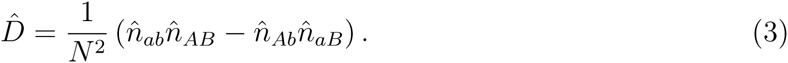

Before the environmental change, the population is well-adapted to its environment and the population size is constant, *N* (*t*) = *N*_0_. The numbers of the respective genotypes in the new generation are determined by multinomial sampling of *N*_0_ individuals, where the probability to sample an individual of type *i* (*i* ∈ {*ab, Ab, aB, AB*}) is given by

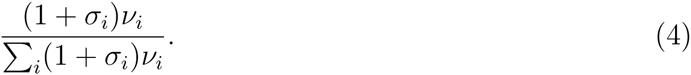

The selection coefficients *σ_i_* quantify selection before the environmental change. We set *σ_ab_* = 0 and assume that all mutants are deleterious relative to the wildtype, *σ_aB_*, *σ_Ab_*, *σ_AB_ <* 0. After the switch in the environment, the population size is variable and will usually initially decline. Individuals of type *i* survive with probability (1 + *s_i_*)*/X* such that their number after selection is Poisson distributed with parameter

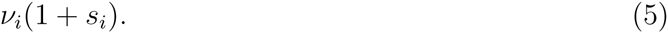

The *s_i_* parametrize the expected growth (for *s_i_* > 0) or decline (*s_i_* < 0) of a type-*i* population after the environmental change. Usually, we assume that only the rescue type can grow under these conditions (*s_AB_* > 0), and *s_ab_*, *s_Ab_*, *s_aB_* are all *<* 0. Epistasis before and after the environmental change is measured as the deviation of fitnesses from multiplicativity, i.e.,

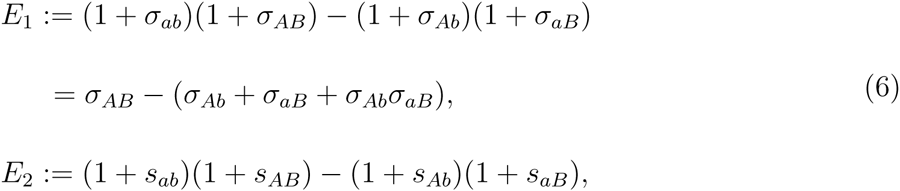

respectively (see Kouyos *et al.* (2007) for the definition of epistasis in discrete vs continuous time models). If the total number of individuals after selection is larger than *N*_0_, it is reduced to *N*_0_. Density regulation hence occurs via a hard carrying capacity, and there is no density dependence for *N* ≤ *N*_0_.

The simulations follow this scheme. We start with a population that is entirely composed of wildtype individuals and let it evolve for 10^4^ generations such that mutation-selection balance is reached before the environment changes (increasing the number of generations did not influence the results). We follow the population dynamics after the environmental change either until the population has gone extinct or until the number of *AB* mutants has grown to 90% carrying capacity and the population can be considered as rescued (in a few cases, we modify the criterion for “rescue” to reduce the simulation time; this is then explicitely stated). The simulation code is written in the *C* programming language, making use of the *Gnu Scientific Library* (Galassi *et al.*, 2009).

## Analysis and Results

Our analytical approach to estimate the rescue probability combines deterministic and stochastic aspects. We focus on populations that are intially large and (mostly) describe the dynamics of the wildtype and the single mutants deterministically. Even if the initial population size is large, however, the number of rescue type individuals (*AB* double mutants) is potentially small and subject to strong stochasticity. This stochastic dynamics depends on all genotype frequencies in the population, which typically change over time in response to environmental change and selection. We address the stochasticity in the number of *AB* mutants by means of branching process theory. The basic mathematical ingredients used are summarized in Appendix S1; Appendix S2 and Appendix S3 contain the derivations of our main results. In Appendix S4, we briefly test the limits of our approximations. Since selection is potentially strong, details of the life cycle need to be taken into account in order to arrive at quantitatively accurate analytical predictions. We take care of these details in the Appendix but neglect them in the main text below, where we summarize our main results.

The probability of evolutionary rescue depends on two factors: the number of rescue types that are generated and their establishment probability in the population after the environmental change. Both quantities are affected by recombination. Mutant genotypes can either be present in the population prior to the switch in the environment or newly arise during population decline. Double mutants, in particular, can either be generated by mutation from single mutants with a constant probability per individual or by recombination of two single mutants with a probability that depends on the (time-dependent) genotype composition in the population. Which route of rescue is most relevant depends on the model parameters for mutation, selection, recombination, and drift. In this section, we progressively describe all routes to rescue. We start with a scenario where single mutants are lethal in the new environment. In that case, rescue entirely relies on double mutants from the standing genetic variation (we assume throughout the analysis that the mutation probability is so small that a direct transition from the wildtype to the double mutant can be neglected). Next, we assume that one of the single mutants is sufficiently viable in the new environment that it can still generate the rescue type by mutation after the environmental change. In both of these scenarios, recombination can be beneficial or detrimental in the old environment due to its effect on the number of rescue genotypes in the standing genetic variation; its effect after the change in the environment is, however, always detrimental (recombination with the wildtype reduces the establishment probability of the rescue type). This is different if both single mutants are viable under the new conditions; in that case, recombination increases the rate at which the rescue type is generated after the environmental change, and – depending on the fitness scheme – this can outweigh the negative effect of recombination. Finally, if single mutants have a fitness larger than one, formation of the double mutant is not required for rescue. We briefly discuss when it is appropriate to neglect its generation in this case. The fitness schemes used in the following four sections are summarized in Table 1.

**Table I.** Fitness schemes used in “Analysis and Results”.

|  | scheme 1 | scheme 2 | scheme 3 |  | scheme 4 |
| --- | --- | --- | --- | --- | --- |
|  |  |  | scenario 1 | scenario 2 |  |
| **ab** | $s_{ab} < 0$ | $s_{ab} < 0$ | lethal | $s_{ab} \leq s < 0$ | lethal |
| **Ab** | lethal | $s_{Ab} < 0$ | $s_{Ab} = s < 0$ | $s_{Ab} = s < 0$ | $s_{Ab} = s > 0$ |
| **aB** | lethal | lethal | $s_{aB} = s < 0$ | $s_{aB} = s < 0$ | $s_{aB} = s > 0$ |
| **AB** | $s_{AB} > 0$ | $s_{AB} > 0$ | $s_{AB} > 0$ | $s_{AB} > 0$ | $s_{AB} > s$ |

## Scheme 1: Single mutants are lethal in the new environment

Within our first scheme, we assume that single mutants are lethal in the new environment (*s_Ab_* = *s_aB_* = -1). This means that de-novo generation of the rescue type is prevented after the change in the environment and rescue – if it happens – happens from double mutants in the standing genetic variation.

Before the change in the environment, single mutants segregate in the population at mutation-selection balance, which we approximate deterministically as constant in time, 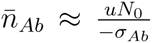, 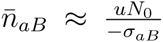 (ignoring the influence of recombination on the frequency of single mutants). Double mutants are generated from single mutants by either recombination or mutation, at a total rate of of 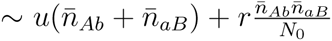. In the presence of wildtype individuals, the double mutant suffers from recombination with the wildtype and gets broken up at rate 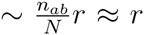 leading to an “effective fitness” of ≈ 1 + *σ_AB_* – *r <* 1. Using this, we can describe the number of *AB* mutants in the standing genetic variation, *n_AB_* by a subcritical branching process with immigration (see Appendix S1.2).

The establishment probability of the rescue type after the environmental change depends on the dynamics of the wildtype. If the wildtype is lethal (*s_ab_* = -1), *AB* mutants are not broken up by recombination and establish with probability 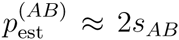 (Haldane, 1927). If the wildtype disappears slowly, with expected extinction time much larger than the typical establishment time of a rescue type, the growth parameter of a rare rescue type is ∼ *s_AB_* – *r* and we can approximate 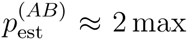[(*s_AB_* − 1)]. In an intermediate parameter range, where the establishment time of the rescue type and the extinction time of the wildtype cannot be separated, we need a more refined approximation (see Appendix S1.2, Eq. (S1.29)): We derive this approximation by treating the wildtype extinction time as a stochastic variable, whose dis-tribution can be estimated. Conditioned on this extinction time, the establishment probability of the rescue type follows from a time-dependent branching process (Uecker and Hermisson, 2011). Finally, for a given *n_AB_* at the time of environmental change, the probability of evolutionary rescue follows as

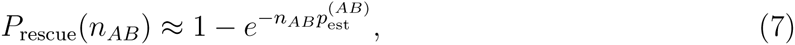

which needs to be averaged over the distribution of *n_AB_*.

Fig. 1 shows the probability of evolutionary rescue for the three possible fitness schemes prior to the environmental change: no epistasis, negative epistasis, and positive epistasis. After the environmental change, epistasis is positive (or zero) since single mutants are lethal. Panels A–C show the behavior for a very large population size (*N*_0_ = 10^8^), where all genotype frequencies prior to the environmental change are close to deterministic; in Panels D–F, the population size is two orders of magnitude smaller (*N*_0_ = 10^6^), and stochasticity in the number of double mutants becomes important (see below).

**Fig. 1:**
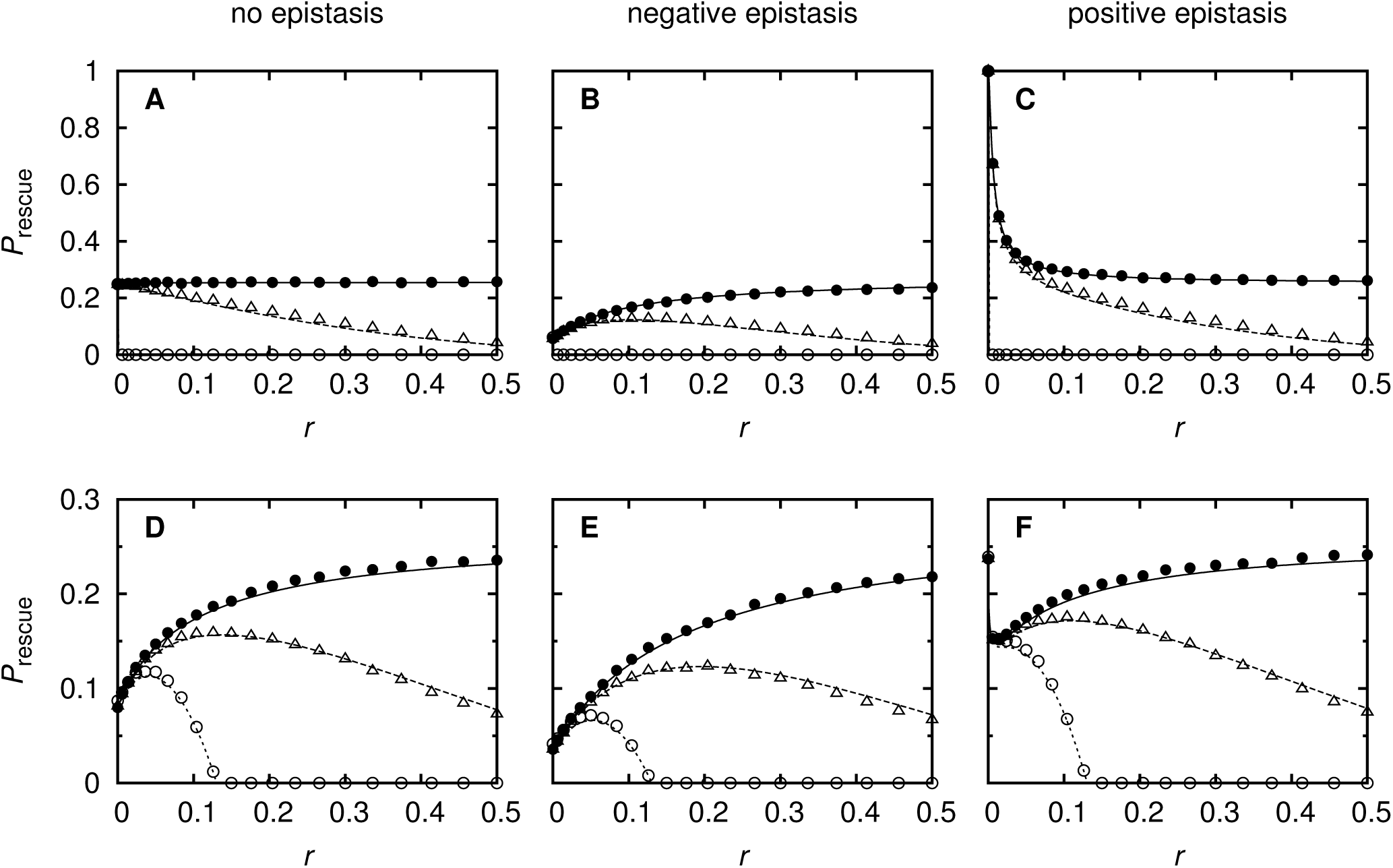
Probability of evolutionary rescue as a function of recombination when single mutants are lethal in the new environment. Filled circles correspond to an instantaneous elimination of the wildtype (*s_ab_* = –1), triangles to an extremely fast (*s_ab_* = –0.99) and empty circles to a slow (*s_ab_* = –0.005) decay of the wildtype population size. Before the switch in the environment, selection against the single mutants is *σ_Ab_* = *σ_aB_* = –0.01, and epistasis is absent (Panels A+D; *σ_AB_* = –0.0199, i.e. *E*_1_ = 0), negative (Panels B+E; *σ_AB_* = –0.1, i.e. *E*_1_ ≈ –0.08), and positive (Panels C+F; *σ_AB_* = –0.0001, i.e. *E*_1_ ≈ 0.02). The other parameter values are: *u* = 10^−5^; *s_AB_* = 0.0015 and *N*_0_ = 10^8^ (Panels A–C); *s_AB_* = 0.15 and *N*_0_ = 10^6^ (Panels D–F). The theoretical curves are based on Eq. (S3.1) (for *s_ab_* = –1), Eq. (S3.10) (for *s_ab_* = –0.005), and Eq. (S3.12) (for *s_ab_* = –0.99). Symbols denote simulation results. Each simulation point is the average of 10^5^ replicates.

If the wildtype is lethal in the new environment (filled circles), recombination affects the probability of rescue only via its effect on the standing genetic variation, and it is instructive to consider this case first. In Appendix S3.1, we derive an approximation for rescue if we can treat all genotype frequencies in the standing genetic variation deterministically (see Eq. (S3.4)); with *σ_Ab_* = *σ_aB_* = *σ* and *σ_AB_* = (*σ_Ab_* +*σ_aB_* +*σ_Ab_σ_aB_*)+*E*_1_, and assuming that all selection coefficients are small, we obtain:

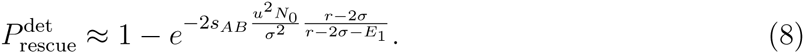

From this, we can read off that for *E*_1_ = 0 (no epistasis, Fig. 1A), the rescue probability is independent of recombination; for *E*_1_ *<* 0 (negative epistasis, Fig. 1B), it increases with *r*; and for *E*_1_ > 0 (positive epistasis, Fig. 1C), it decreases with *r*. For *r* = 0, the rescue probability depends strongly on epistasis 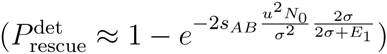, while for *r* ≫ max [|2*σ*|, |2*σ* + *E*_1_|], the rescue probability becomes independent of epistasis 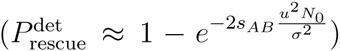. This is expected from deterministic theory: with no epistasis, the population is in linkage equilibrium and recombination has no effect. Negative epistasis leads to negative linkage disequilibrium (LD) and recombination is favorable since it increases the number of *AB* mutants. Vice versa, positive epistasis leads to positive linkage disequilibrium and recombination is deleterious since it decreases the number of *AB* mutants (Felsenstein, 1965). For sufficiently strong recombination, the population approaches linkage equilibrium irrespective of epistasis.

In Panels D–F, population size is smaller by two orders of magnitude. The growth rate of the rescue genotype is two orders of magnitude larger such that *s_AB_N*_0_ is the same as in Panels A–C. The deterministic prediction for the rescue probability Eq. (8) is hence unchanged. However, although the population size before the decline contains 10^6^ individuals, stochastic fluctuations in the number of double mutants *n_AB_* become important in this regime. Symmetric fluctuations in *n_AB_* do not have symmetric effects on rescue, since the rescue probability (Eq. (7)) is a concave function of *n_AB_* (it does not help getting rescued twice). Negative fluctuations thus have a stronger effect on *P*_rescue_ than positive fluctuations and drift reduces the probability of evolutionary rescue. This effect is most pronounced for tight linkage, but is attenuated by recombination (e.g. Panel D). Since recombination pulls genotype frequencies closer to linkage equilibrium, it overall dampens fluctuations in LD and along with it fluctuations in *n_AB_*. For positive epistasis prior to the environmental change (Panel F), this results in a non-monotonic dependence of *P*_rescue_ on recombination. For small *r* (*r* ≲ 2|*σ*|, see Eq. (8)), recombination is deleterious since it breaks up the positive LD built up by epistasis; for larger *r*, the positive effect of recombination (attenuating drift) dominates. Fig. S3.1 in Appendix S3 disentangles the effects of epistasis and drift.

If the wildtype is not lethal in the new environment (open symbols), recombination is deleterious after the environmental change. Note that even a slight increase of the wildtype fitness above lethality drastically influences the outcome for recombination *r*≳*s_AB_* (empty triangles). The presence of the wildtype during the first few generations after the environmental change is sufficient to break up double mutants and to hamper their establishment probability significantly. If the wildtype is quite fit (open circles) and recombination strong (*r*≳*s_AB_*), rescue becomes impossible. This corresponds to observations in populations of constant size where the crossing of fitness valleys is prevented by strong recombination (Crow and Kimura, 1965; Jain, 2010; Weissman *et al.*, 2010).

For a slowly decaying wildtype, the results are robust to deviations of the fitness of single mutants from strict lethality; however, for a lethal wildtype, chances of rescue increase considerably when single mutants have a fitness slightly larger than zero (see Appendix S3.1 and Fig. S3.2; see below for the scenario *s_Ab_* = *s_aB_* = *s* and *s_ab_* = -1).

## Scheme 2: One single mutant is viable, the other is lethal

Viability of one of the single mutants, say *Ab*, opens up new pathways to rescue since new double mutants can be generated by mutation after the switch in the environment. Rescue can occur via (a) double mutants from the standing genetic variation as in the previous paragraph, mutation of single mutants from the standing genetic variation after the environmental change, and – if the wildtype is viable in the new environment – (c) two-step mutation after the environmental change (i.e., generation of single mutants and subsequently double mutants, both by de-novo mutation). Our aim in this section is to study the relative importance of standing variation vs new mutation in two-locus rescue. In the Appendix, treat both the *Ab* single mutants and the *AB* double mutants stochastically, using a two-type branching process, to derive the rescue probability. This is necessary for a quantitatively precise approximation. For a qualitative assessment of the relative importance of rescue pathways, it is sufficient to stick to a simple deterministic treatment of all single mutant dynamics.

If the wildtype is either lethal or disappears sufficiently slowly, each single mutant has approximately the same chance to generate a permanent lineage of *AB* mutants, independently of whether it is already present at the time of environmental change or arises later. In order to compare the relative importance of pathways (b) and (c), it is hence sufficient to compare the number of *Ab* mutants in the standing genetic variation 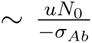 with the number of *Ab* mutants that get newly generated during population decline 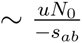 (assuming 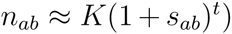). Accordingly, the contribution from pathway (c) is larger than that from (b) if *s_ab_ = σ_Ab_*. In order to compare pathways (a) and (b), we compare the exponent of Eq. (8), assuming *σ_Ab_* = *σ_aB_*, with the number of successful *AB* mutants issued from *Ab* mutants in the standing genetic variation. If we assume a deterministic decay of the *Ab* mutants in the new environment, 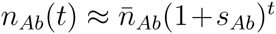 the latter is given by *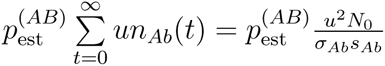*, where 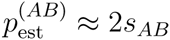 if the wildtype is lethal and 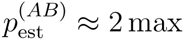 if the wildtype disappears slowly. With *E*_1_ = 0 or *r* large, this contribution is larger than rescue via pathway (a) if *s_Ab_ > σ_Ab_*, i.e., if the growth parameter of single mutants is larger in the new than in the old environment; for *r* → 0, it is larger if *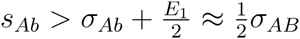*. Overall, we obtain for rescue via pathway (b) or (c):

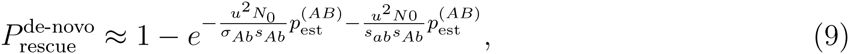

where the first summand in the exponent accounts for pathway (b) and the second summand for pathway (c), using that every single mutant leaves on average 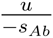 double mutant offspring. If the wildtype is lethal (Fig. 2A), the contribution of route (b) to rescue is independent of recombination, and recombination has an influence on rescue only via its effect on the number of double mutants in the standing genetic variation (compare Fig. 2A with Fig. 1D). If the wildtype is viable (Fig. 2B), recombination is deleterious after the environmental change (cf. Fig. 1).

**Fig. 2:**
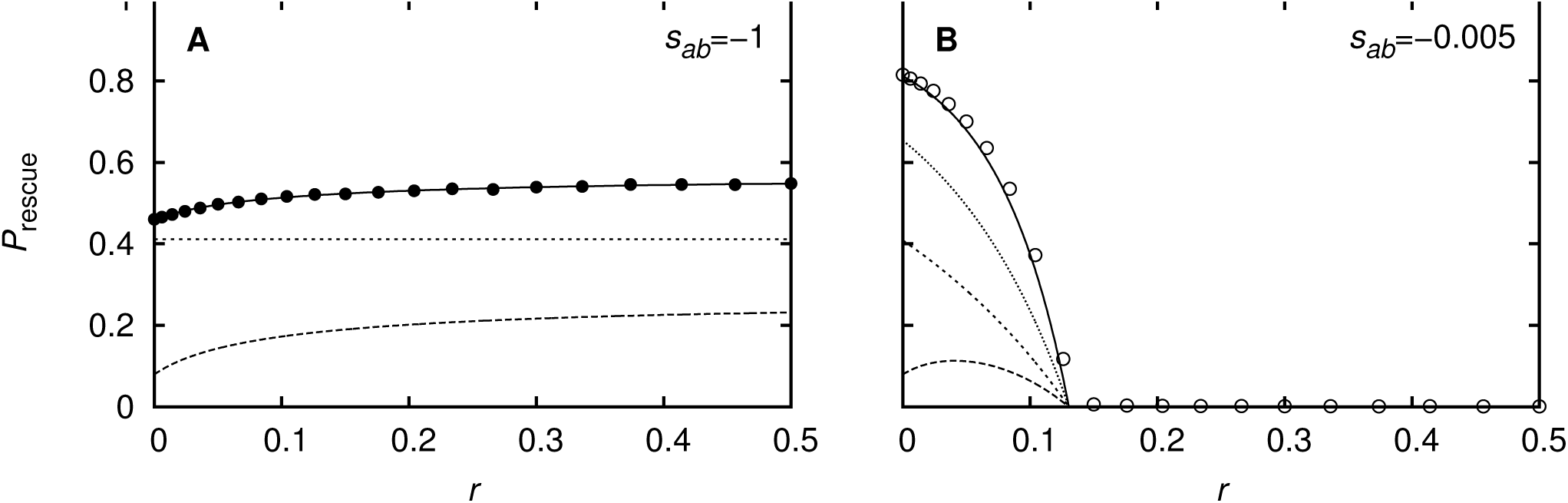
Probability of evolutionary rescue as a function of recombination when one single mutant is lethal and the other one viable. In Panel A, the wildtype is lethal; in Panel B, it is only mildly deleterious. Solid line (with simulation dots): total probability of evolutionary rescue; long-dashed line: rescue only via double mutants from the standing genetic variation; short-dashed line: rescue via single mutants from the standing genetic variation (which subsequently mutate); dotted line (only Panel B): rescue from two-step de-novo mutation. Parameter values: *σ_Ab_* = *σ_aB_* = –0.01, *σ_AB_* = –0.0199 (i.e., no epistasis before the environmental change), *s_Ab_* = –0.005, *s_aB_* = –1, *s_AB_* = 0.15, *u* = 10^−5^, *N*_0_ = 10^6^. The theoretical curves are based on Eq. (S3.14) (Panel A) and Eq. (S3.20) (Panel B). Each simulation point is the average of 10^5^ replicates.

The more precise analysis in Appendix S3.2 extends to *s_Ab_* = 0. Then, rescue does not require the generation of the double mutant. We discuss in the Appendix, when focussing on establishment of type *Ab* is sufficient under these conditions.

## Scheme 3: Both single mutants are viable in the new environment

In our third scheme, we turn to scenarios in which single mutants have absolute fitness 0 *<* 1 + *s_Ab_*, 1 + *s_aB_ <* 1 in the new environment such that the last pathway to rescue opens up: in addition to the previous routes, the rescue mutant can now be generated by recombination after the environmental change. As in the phase prior to the environmental switch, the net role of recombination after the environmental change depends on the linkage disequilibrium among types: with negative LD, the net effect of recombination is to generate rescue mutants, with positive LD, it rather breaks them up. The expected LD, in turn, depends on the growth rates (fitnesses) of the four types: positive/negative epistasis entails positive/negative LD. With a switch in the selection regime at the time of environmental degradation, we can thus distinguish four basic scenarios, combining positive or negative epistasis before the switch with either positive or negative epistasis after the change, keeping or reversing the role of recombination.

For our analysis, we consider two cases for the fitness scheme after the environmental change: (1) negative epistasis with 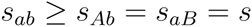 and *s_ab_* = -1 (i.e., the wildtype is lethal) and (2) positive epistasis with 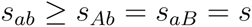. Epistasis in the original environment is positive or negative. For simplicity, we assume equal single-mutant fitnesses, *σ_Ab_* = *σ_aB_* = *σ*. Note that, with this choice, we have *n_Ab_*(*t*) = *n_aB_*(*t*) for all times, as long as drift can be ignored.

Rescue from double mutants in the standing genetic variation can be evaluated as above (Scheme 1) with 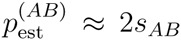 for scenario 1 and *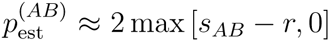* for scenario 2. In order to determine the total probability of evolutionary rescue, we need to add rescue from double mutants that originate after the environmental change. These are generated at a time-dependent rate 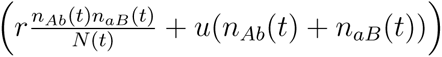.In scenario (1), with *n_Ab_*(*t*) = *n_aB_*(*t*) and *N* (*t*) = *n_Ab_*(*t*) + *n_aB_*(*t*) (which holds as long as the *AB* mutant is rare), this simplifies to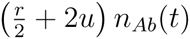. The rate of decline of single mutants is considerably enhanced by recombination in this scenario, since half of all recombination events occur among *Ab* and *aB* types. Each such recombination event breaks both single mutants up. The single mutant types hence decay at rate 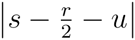, generating approximately 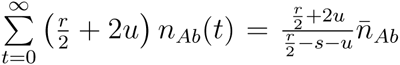 double mutants on their way to extinction. Each of these double mutants establishes a permanent lineage with probability 2*s_AB_*. The combination of these two rescue pathways – generation of the rescue type by mutation or recombination after the environmental change – is hence given by

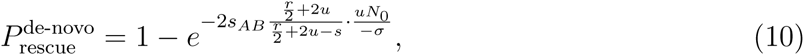

which increases with *r*. If epistasis is negative prior to the environmental change, recombination is hence advantageous in both phases and *P*_rescue_ increases with *r* (Fig. 3A). If, on the other hand, epistasis is positive in the old environment, the effect of recombination changes from negative to positive between the two phases. The negative effect in the old environment and the positive effect in the new environment act on different recombination scales: the negative effect levels off for *r* ≫ 2|*σ*| (see Eq. (8)). As can be seen from Eq. (10), the positive effect of recombination levels off once *r* ≫ –*s*. In Fig. 3B, selection is stronger in the new environment (|*s*| ≪ |*σ*). Moreover, 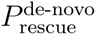 is small for weak recombination, since the single mutants decay rapidly after the environmental change. Therefore, the negative effect of recombination dominates for small *r*; the positive effect takes over as recombination increases.

**Fig. 3:**
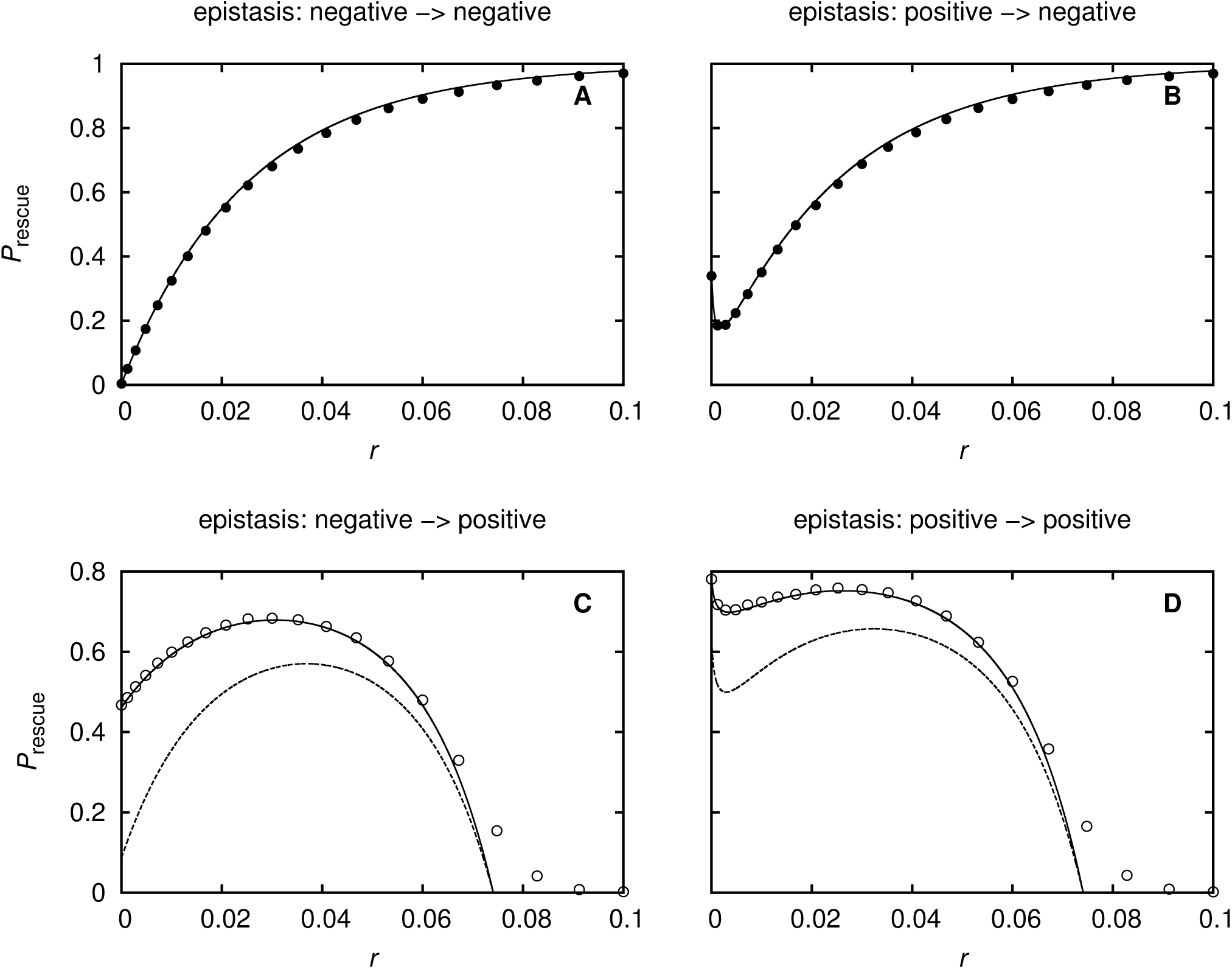
Probability of evolutionary rescue as a function of recombination for different patterns of epistasis before and after the environmental change. Solid line: total probability of rescue; dashed line: probability of rescue from the standing genetic variation (i.e., without new mutations after the environmental change). In Panels A and B, both lines coincide. The analytical predictions in Panels A and B are based on Eq. (S3.32) and its components; the analytical predictions in Panels C and D are based on Eq. (S3.38) and its components. Parameter values: *N*_0_ = 10^8^, *u* = 2 · 10^−6^, *σ_Ab_* = *σ_aB_* = –0.01; first row (A+B): *s_Ab_* = *s_aB_* = –0.5, *s_ab_* = –1, *s_AB_* = 0.002, and *σ_AB_* = –0.1 (i.e., *E*_1_ ≈ –0.08, Panel A), *σ_AB_* = –0.0001 (i.e., *E*_1_ ≈ 0.02, Panel B); second row (C+D): *s_ab_* = *s_Ab_* = *s_aB_* = –0.03, *s_AB_* = 0.08, and *σ_AB_* = –0.1 (i.e., *E*_1_ ≈ –0.08, Panel C), *σ_AB_* = –0.0001 (i.e., *E*_1_ ≈ 0.02, Panel D). Symbols denote simulation results. Each simulation point is the average of 10^5^ replicates.

Scenario (2), used in Fig. 3C+D, is more complex. The proportion of single mutants changes during population decline (even for *s_ab_* = *s*, since new single mutants arise during population decline). The rate at which single mutants recombine to generate double mutants is hence not constant and the approximation for the total number of double mutants that get newly generated after the environmental change does not take a simple form. However, it is still possible to calculate it analytically, see Eq. (S3.37) and (S3.43a) in S3. Due to the presence of the wildtype, the rate is much smaller than 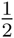. Moreover, recombination reduces the establishment probability of the rescue mutant to 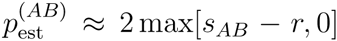, rendering rescue impossible for strong recombination. If epistasis is negative prior to the environmental change such that *AB* mutants are underrepresented in the standing genetic variation without the aid of recombination, we find that *P*_rescue_(*r*) displays an intermediate maximum (Fig. 3C, and Fig. S3.3). If epistasis is positive both in the old and in the new environment, deterministic theory predicts that recombination is always harmful. However, in finite populations with a small number of double mutants, recombination has a positive effect by attenuating the effect of drift (as described for Scheme 1, Fig. 1). As a consequence, *P*_rescue_ displays a minimum and a maximum in Panel D of Fig. 3.

## Scheme 4: Both single mutants have fitness greater than one

If single mutants have fitness greater than one, formation of the double mutant is not required for rescue, but can still increase the rate of rescue if the double mutant is considerably fitter than the single mutants. However, formation of the double mutant comes at the cost of losing two single mutants. Keeping the single mutants intact can therefore be better for rescue than generating the double mutant if the latter is only slightly fitter than the single mutants.

For simplicity, we consider scenario 1 from the previous section with lethal wildtype after the change but allow *s_Ab_* = *s_aB_* = *s* to be greater than zero. Under these conditions, recombination cannot break the double mutant after the change in the environment. The role of recombination simply is to convert the different rescue types into each other, more precisely to turn two single mutants (that are now also rescue genotypes) into one double mutant. One individual of type *AB* establishes a permanent lineage with probability ≈ 2*s_AB_* whilst one individual of type *Ab* (or *aB*) establishes a permanent lineage of single mutants with probability 2*s*. Intuitively, conversion of two single mutants into one double mutants therefore pays off if *s_AB_* ≫ 2*s* and is deleterious for rescue if *s_AB_* ≪ 2*s*. Recombination hence increases the rate of rescue if *s_AB_* ≫ 2*s* and decreases the rate of rescue if *s_AB_* ≪ 2*s*; it has little effect if *s_AB_* ≈ 2*s* (see Fig. 4). We formalize this argument in Appendix S3.4.

**Fig. 4:**
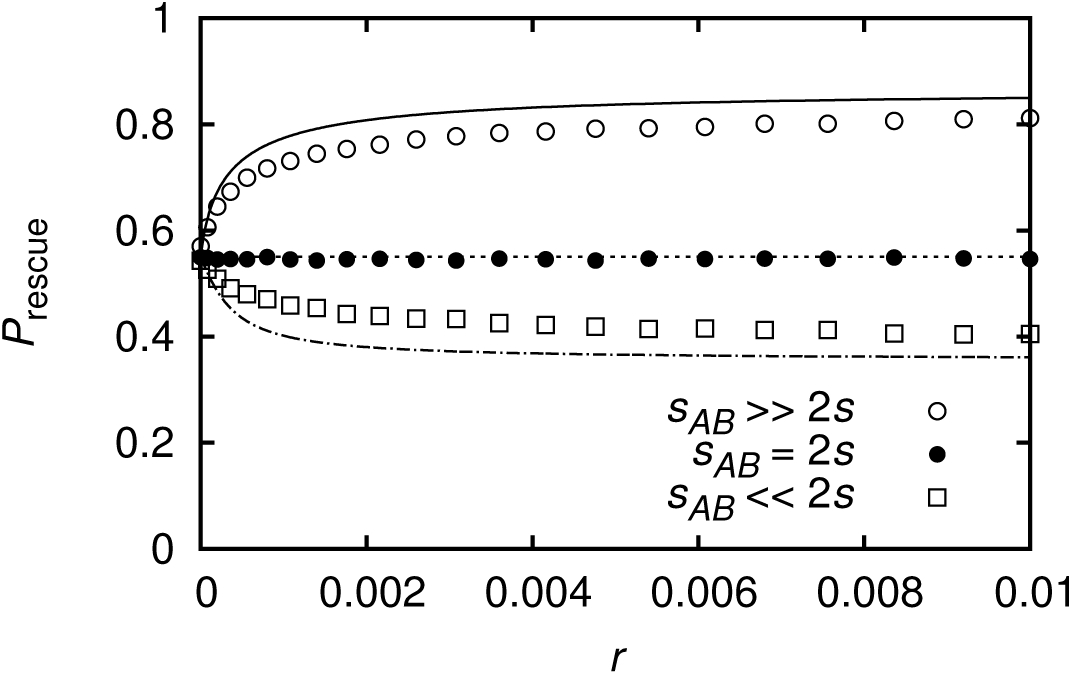
Probability of evolutionary rescue as a function of recombination. For *s_AB_* ≫ 2*s*, *P*_rescue_ increases with recombination; for *s_AB_* = 2*s*, recombination has no effect on rescue; for *s_AB_* ≪ 2*s*, *P*_rescue_ decreases with recombination. Parameter values: *N*_0_ = 10^7^, *u* = 2 · 10^−6^, *σ_Ab_* = *σ_aB_* = –0.01, *σ_AB_* = –0.0199, *s_ab_* = –1, *s_Ab_* = *s_aB_* = 10^−4^; empty circles: *s_AB_* = 5 · 10^−4^ ≫ 2*s*; filled circles: *s_AB_* = 2 · 10^−4^ = 2*s*; squares: *s_AB_* = 0.11 · 10^−4^ ≪ 2*s*. The theoretical predictions are based on Eq. (S3.45) combined with Eq. (S3.1). Symbols denote simulation results. For the simulations, we consider the population as rescued if the total number of mutant genotypes has reached 20% carrying capacity. Each simulation point is the average of 10^5^ replicates.

## Notable observations

To conclude, we point out two effects of recombination on rescue probabilities which might contradict spontaneous intuition. First, with recombination, a high frequency of wildtype individuals after the environmental change is a potent force to inhibit rescue by double mutants. Consequently, a slower decay of the wildtype population often reduces, rather than promotes, the chances for population survival. While a slower decay leads to a higher rate of new single mutants, the latter are less likely to meet and recombine in the presence of a dominating wildtype. Also, if a double mutant is generated, it is likely to be broken up by recombination. Depending on the strength of recombination, the rate of rescue decreases or increases or displays an intermediate minimum as a function of the wildtype fitness (see Fig. 5A; cf. also Fig. S3.3). Indeed, we obtain a clear decrease in *P*_rescue_ for almost the entire range of wildtype fitness *s_ab_*, unless recombination is extremely weak. Only for very high wildtype fitness, approaching viability (*s_ab_* = 1), *P*_rescue_ steeply increases again (cf. Fig. 5A).

**Fig. 5:**
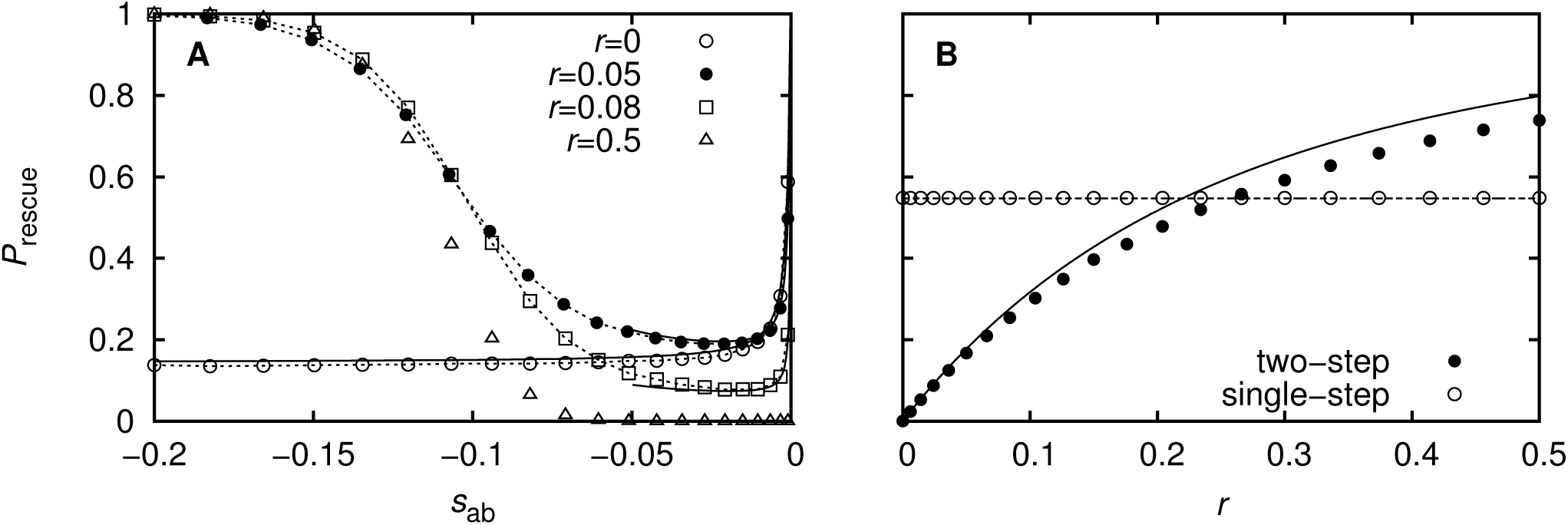
Probability of evolutionary rescue. Panel A: Probability of evolutionary rescue as a function of wildtype fitness for various values of *r*. Solid curves consitute analytical predictions and are based on Eq. (S2.2) (*r* = 0) and Eq. (S3.38) with Eq. (S3.43a) (*r =* 0). Dotted lines connect simulation points and are included to guide the eye. Parameter values: *σ_Ab_* = *σ_aB_* = –0.01, *σ_AB_* = –0.0199, *s_Ab_* = *s_aB_* = –0.05, *u* = 10^−5^, *N*_0_ = 10^6^, *s_AB_* = 0.1. Panel B: Probability of evolutionary when the rescue mutant is one or two mutational steps away. The analytical predictions are based on Eq. (S3.32) (for two-step rescue) and Eq. (S3.49) (for single-step rescue). Parameter values: *u* = 2 · 10^−6^, *N*_0_ = 10^7^, *σ_Ab_* = *σ_aB_* = –0.01, *σ_AB_* = –0.1, *s_ab_* = –1, *s_Ab_* = *s_aB_* = –0.5, *s_AB_* = 0.002. Symbols denote simulation results. For Panel A, we consider a population as rescued when the number of double mutants has reached 20% carrying capacity (increasing the threshold to 30% did not change the results). Each simulation point is the average of 10^5^ replicates.

Second, with recombination, single mutant types can act as an important buffer to environmental change, even if they are not able to rescue a population on their own. As a consequence, two-step rescue does not need to be less likely than single-step rescue with a single locus and direct generation of the rescue type from the wildtype by a single mutation (see Fig. 5B). Imagine a situation where the wildtype is lethal in the new environment. Single-step rescue now relies entirely on the rescue type individuals that are present at the time of environmental change. In two-step rescue, single mutants might still be present in the new environment and can generate the rescue mutant by mutation or recombination at a high rate. Even the number of double mutants in the standing genetic variation can be higher for two-step than for single-step rescue if the mutation rate is high or epistasis strongly positive (see Appendix S3.5 for more details).

## Discussion

Following severe environmental change, populations might find themselves maladapted to the new conditions, and a race between population decline and adaptive evolution begins. In conservation biology, the desired outcome of this race is persistence of the population; in medicine, in contrast, one aims at eradication of the pathogen from the human body. Adaptation to the new environmental conditions is often contingent on allelic changes at more than one locus. This holds, in particular, for resistance to multiple drugs in combination drug therapy or pesticide mixtures in agriculture. Complex rescue, requiring adaptation at multiple loci, is expected to lead to severely reduced probabilities of rescue (or resistance). However, the prediction of these probabilities can be surprisingly complicated if there is recombination among the target loci.

In this paper, we have analyzed a generic two-locus model in order to clarify the role of recombination in evolutionary rescue. We find that depending on the fitness scheme of mutations, recombination can make two-step rescue even more likely to happen than single-step rescue but it can also prevent rescue entirely (see Fig. 3C+D, and Fig. 5B). Recombination acts to reduce positive or negative LD that is build up by epistasis and it weakens fluctuations in LD caused by genetic drift. Since there are two phases of selection – before and after the environmental change – and drift, recombination acts threefold, where the effects can go in different directions (increasing or decreasing the rate of rescue) and show at different scales. As a consequence, dependence of rescue on recombination can be non-monotonic with multiple ups and downs (see e.g. Fig. 3D). Also the dependence of rescue on the wildtype dynamics is non-trivial, with a slow decline of the wildtype not always being better for population survival than a fast eradication (see Fig. 5A).

### The role of epistasis

As well-known from classical population genetics, whether recombination speeds up or slows down adaptation strongly depends on the sign of epistasis (Felsenstein, 1965). In scenarios of evolutionary rescue, there are two phases of selection with potentially epistatic interactions between loci, before and after the change in the environment, and the role of recombination is affected by epistasis in both phases. Experimentally, the strength and sign of epistasis have been measured across a variety of systems (Bonhoeffer *et al.*, 2004; Kouyos *et al.*, 2007; Trindade *et al.*, 2009; Silva *et al.*, 2011), reporting all forms of epistasis. Moreover, several studies have investigated the influence of the environment on the epistatic interactions between mutations, finding that both the strength and the sign of epistasis can change as the environment changes (Remold and Lenski, 2004; Lalic and Elena, 2012; De Vos *et al.*, 2013; Flynn *et al.*, 2103). This shows that for a comprehensive picture of the role of epistasis on adaptation upon environmental change, all possible fitness schemes in both environments need to be studied.

Epistasis leads to linkage disequilibrium which is broken up by recombination. Recombination thus counteracts selection. The scale at which recombination is effective is determined by the strength of epistatic selection. Since the strength of selection can be different before and after the environmental change, the relevant recombination scales can be different, too, and if the sign of epistasis changes with the environement, the probability of evolutionary rescue depends non-monotonically on recombination.

We can compare our results with classical models for the crossing of fitness valleys in populations of constant size. In these models, a small but non-zero recombination rate minimizes the time to get from one fitness peak to another, while strong recombination hampers or even prevents the crossing of valley in large populations (Jain, 2010; Weissman *et al.*, 2010; Altland *et al.*, 2011). Epistasis is positive in this scenario. However, since double mutants are initially absent, LD is negative during a first transient phase. As the frequency of double mutants increases (supported by recombination), LD turns positive and recombination counteracts any further increase of double mutants. The valley-crossing scenario thus compares to a rescue situation with negative epistasis in the old and positive epistasis in the new environment. Indee, we obtain analogous results under these conditions (see Fig. 3C).

### The role of drift

We find that genetic drift has a strong influence on rescue probabilities, even in very large populations (see e.g. Fig. S2.1 with *N*_0_ = 10^8^). This may seem surprising, but can be understood because the decisive quantity for rescue is the number of double mutants, which is potentially small even if the total population size is large. Importantly, stochastic fluctuations in the number of rescue types do not only entail a stochastic outcome (extinction or survival), but also have a directional (negative) effect on the rescue probability. This is because for any given population, the probability of evolutionary rescue is a concave function of the number of rescue mutants in the standing genetic variation. Consequently, the decrease in the rescue probability due to negative fluctuations in the number of rescue types is larger than the corresponding increase due to positive fluctuations. This effect is strong if the number of double mutants is small and their establishment probability large.

The effect is most prominent for two-step adaptation at a single locus, i.e., in the absence of recombination. Previous theory for two-step rescue for that case has described all genotype frequencies in the standing genetic variation deterministically (Iwasa *et al.*, 2003, 2004). While this is appropriate when the number of double mutants is large and their establishment probability small, the approach strongly overestimates the probability of evolutionary rescue if these conditions are not met (see S2 and Fig. S2.1).

Recombination attenuates this effect of drift by pulling the number of double mutants closer to its expected value and thus increases the probability of rescue. We find that the decrease of fluctuations in LD (and in the number of double mutants) affects rescue equally or sometimes even more strongly than the reduction of directional LD (increasing or decreasing the mean number of double mutants) which has been built up by epistasis. We finally note that the interaction of drift and recombination described here is different from the effect of recombination in the presence of Hill-Robertson interference in finite populations that has been described previously (Barton and Otto, 2005; Roze, 2014). This latter mechanism works through a small bias towards negative LD on average because selection acts asymmetrically on symmetric fluctuations in LD. This is negligible in our model, while fluctuations in LD turn out to be very prominent.

### The population dynamics

As long as the rescue type is rare, the population dynamics are shaped by the dynamics of the wildtype and the single mutants. Dependence of rescue on the dynamics of single mutants is as expected: the slower the decay, the higher the chance of rescue. The dependence on the dynamics of the wildtype population size is more complex and shaped by two opposing effects. By mutation of wildtype individuals, single mutants arise, which can subsequently mutate or recombine to generate the rescue type. On the other hand, recombination with wildtype individuals breaks the rescue type up. Presence of the wildtype hence increases the rate at which the rescue mutant is generated but decreases its establishment probability. As a consequence, dependence of rescue on the rate of decline of the wildtype population is non-trivial. We find that the rate of rescue decreases with wildtype fitness over a large parameter range but it can also increase and be overall non-monotonic as a function of wildtype fitness (see Fig. 5A). For strong recombination, a high frequency of wildtype individuals prevents rescue entirely. Importantly, dependence of rescue on the presence or absence of the wildtype can be very sensitive such that even a slight increase of the wildtype fitness above lethality can signficantly reduce the probability of population survival (see Fig. 1).

Violations of the simple rule for drug therapy to “hit hard” (the faster the wildtype population disappears, the lower the risk of resistance) have been found before as a consequence of competitive release: if the fitness of the rescue type is density dependent, a fast eradication of the wildtype enhances rescue by freeing up resources (Torella *et al.*, 2010; Read *et al.*, 2011; Peñamiller *et al.*, 2013; Uecker *et al.*, 2014). Note that here, we find that the rule is violated also in a model without competition for resources and with density-independent fitness. Our results imply that in order to prevent resistance, it is of vital importance to suppress the single mutants efficiently while it can be preferable to remove the wildtype slowly. Otherwise the single mutants can act as a reservoir for mutations from which the rescue type can be generated even if the single mutants are not long-term viable themselves and even if they have a very low fitness (see Fig. S3.2). As a consequence, two-step rescue can be even more likely than single-step rescue (see Fig. 5B).

Normally, we expect that in the presence of several drugs, a mutant that is resistant to one of the drugs has a higher fitness than the wildtype strain. For example, Chait *et al.* (2007) find this behavior when they expose wildtype and doxycycline-resistant *E. coli* bacteria to a drug combination of doxycycline and erythomycin. The two drugs act synergistically, i.e., the wildtype has a lower fitness in the presence of both drugs than expected from the single-drug effects. In a combination of doxycycline and ciprofloxacin, however, Chait *et al.* (2007) show that the doxycycline-resistant mutant is even less fit than the wildtype at certain drug concentrations (we apply such a fitness scheme in the limit of lethal single mutant(s) in Fig. 1 and in Fig. 2B). At these concentrations, the two drugs display “suppression interaction”, i.e., the wildtype has a higher fitness in the presence of both drugs than in the presence of just one drug, which is an extreme form of antagonistic drug interactions (one drug attenuates the effect of the other). Based on these findings, Torella *et al.* (2010) developed a mathematical model for the evolution of multi-drug resistance under synergistic and antagonistic drug interactions (implementing no form of recombination). The model shows that resistance evolves less easily under antagonistic interactions but again only if competition among cells is high (for experiments, see Hegreness *et al.* (2008)). Our results suggest that even without competition, antagonistic drug interactions (with a relatively fit wildtype but unfit single mutants) can strongly hamper the evolution of resistance for infections with pathogens that readily recombine in vivo, such as HIV.

### Limitations and extensions

Our analysis gives a comprehensive overview of the role of recombination in the two-locus model for evolutionary rescue. However, quantitatively accurate analytical results are only possible in parts of the parameter range. Most importantly, if both single mutant types are viable and can recombine, we need to describe their frequencies deterministically. This requires a sufficiently large population size. We illustrate the limits of this approach in S4.

Our model describes the most basic situation both on the genetic and on the ecological side (two loci, two alleles per locus, panmictic population, sudden environmental shift). On the genetic side, the incorporation of more loci and the consideration of more complex fitness landscapes constitutes a logical next step. On the ecological side, a variety of extensions would help to gain a more comprehensive understanding of two-step rescue with recombination. A gradual instead of sudden deterioration of the environment influences the population dynamics which, as discussed above, plays a relevant role in rescue. Likewise, population structure with parts of the habitat deteriorating later than others changes the rate of disappearance of the wildtype (Uecker *et al.*, 2014).

Our “minimal model” approach means, of course, that the results cannot be directly applied to concrete cases of resistance evolution. While we expect that the basic principles observed in this study should hold under general conditions, further factors need to be included for specific cases. For example, recombination in HIV is density dependent since multiple infection of a cell is required for recombination to occur. Also, phenotypic mixing does not allow for a simple correspondence between phenotype and genotype and long-lived cells lead to specific population dynamics. Likewise, all three forms of bacterial recombination – conjugation, transduction, transformation – differ significantly from the simple recombination scheme applied in this study, requiring two mating types, the action of bacteriophages, or the release and uptake of DNA molecules into/from the environment.

We entirely focused on the probability of evolutionary rescue, leaving other aspects of rescue unexplored. It would be interesting to find out how recombination affects the time to rescue and the shape of population decline and recovery given rescue occurs (cf. Orr and Unckless (2014) for a study on these aspects in a one-locus model). The mimimum population size of the U-shaped rescue curve is predictive for the amount of standing genetic variation that is preserved over the course of adaptation. The latter in turn affects how well a population can respond to subsequent environmental change.

To summarize, we have analyzed a generic model of two-step rescue with recombination. We find that the role of recombination in rescue is complex and ambivalent, ranging from highly beneficial to highly detrimental. Since recombination of rescue mutants with wildtype individuals destroys the rescue type, a fast eradication of the wildtype can counterintuitively promote rescue even in the absence of competition for resources. A high fitness of single mutants always fosters rescue even if they cannot persist at long-term in the environment themselves. Recombination of single mutants that provide a buffer to environmental change can render two-step rescue even more likely than one-step rescue.

## Acknowledgments

The authors thank Nick Barton, Sally Otto, Denis Roze, and Jitka Polechová for helpful discussions and/or comments on the manuscript. This work was made possible by by a “For Women in Science” fellowship (L’Oréal Ö sterreich in cooperation with the Austrian Commission for UNESCO and the Austrian Academy of Sciences with financial support from the Federal Ministry for Science and Research Austria) and European Research Council Grant 250152 (to Nick Barton).

